## Appendix for "Geometric models for robust encoding of dynamical information into embryonic patterns"

#### 1 A two-enhancer model reproduces dynamical features of Tribolium segmentation

In [1], two of us proposed a model of Tribolium segmentation relying on the interplay of two sets of enhancers. In short, two sets of enhancers (static $S(P)$, dynamic $D(P)$) were used, where the role of parameter g is played by morphogen *Caudal* (*cad*) [2] (Figure 1–figure supplement 2). $S(P)$ encodes a multistable system and $D(P)$ a sequential cascade of genetic expression of gap genes ($hb$, $Kr$, $mlpt$, $gt$). This system was found to implement a "speed gradient" model, where the speed of traveling waves of gap genes from posterior to anterior depended on the level of *cad* concentration (Figure 1–figure supplement 2B-D). This led to robust patterning of the embryo (Figure 1–figure supplement 2E) but the mathematical origin of the speed gradient was not explained.

To better understand the underlying dynamics of the system, we consider the time courses of multiple cells at different positions and thus with different final fates. Figure 1–figure supplement 2F shows the projection of the cells' dynamics on a 2D plane corresponding to the first two genes expressed in the cascade ($Kr$ and $hb$), as well as a typical flow for different values of $cad$ while keeping the other genes ($mlpt$, $gt$ and $X$) at zero. Importantly, for $cad$ = $0.13$, we see the appearance of a new fixed point (green disk on Figure 1–figure supplement 2F).

We make four observations:

- The flow of the system is canalized. The trajectories of the cells stay very close to one another in phase space.
- As *cad* is lowered, the new fixed point appears very close to the common trajectory of all cells (Figure 1–figure supplement 2F, top row), and clearly separates the trajectories of cells ending up at different fates (Figure 1–figure supplement 2F, bottom row).
- When *cad* further decreases, the new fixed point moves in the high $hb$, low $Kr$ region, corresponding to the eventual fate of Cell 1.
- When the new fixed point appears, the flow of cells past this fixed point is slowed down (Figure 1–figure supplement 2G).

These four observations offer a concise explanation to the “speed gradient” model: as the system gets closer to the bifurcation happening at $cad = 0.13$, the system is slowing down because of the future fixed points appearing on the trajectory. Intuitively, this is due to the fact that a fixed point corresponds to a frozen state, and thus to an infinite time-scale (static). When *cad* varies, the system has to interplay between a non-zero time-scale (dynamics) and such infinite time-scale, and it thus makes sense *a priori* that in between, the time-scale of the system diverges. This mechanism is close in principle to the critical timing proposed in [3].

### 2 List of the functions used for the dynamics of each model

#### 2.1 Gene network models

In the gene network models, biochemical interactions between genes are modeled explicitly. Ordinary differential equations (ODEs) represent the dynamics of the concentration of the proteins that are encoded by the genes in the network. The deterministic part of the dynamics is composed of a protein production term and a protein degradation term. The production rate of a given protein can be altered by the interactions between the genes. Hill functions are use to model repression and activation of the production of a given protein by the genes. When multiple genes affect the concentration of a protein, the Hill functions corresponding to each interaction are multiplied. In the simulations, we set to 1 the maximal production rate of all proteins. Similarly, we set the degradation rate of all proteins to 1. In Eq. 1 of the main text, $C(P)$ encodes the degradation term, and $\Theta_S(g)\ S(P) + \Theta_D(g)\ D(P)$ represents the production term.

##### 2.1.1 3-gene models

The proteins associated to the 3 genes are named arbitrarily $A$, $B$ and $C$:

$$P = \begin{bmatrix} A \\ B \\ C \end{bmatrix} \qquad C(P) = \begin{bmatrix} -A \\ -B \\ -C \end{bmatrix} \qquad D(P) = \begin{bmatrix} \frac{1}{1+\left(B/K_D^{B\dashv A}\right)^5} \\ \frac{1}{1+\left(C/K_D^{C\dashv B}\right)^5} \\ \frac{1}{1+\left(A/K_D^{A\dashv C}\right)^5} \end{bmatrix} \qquad S(P) = \begin{bmatrix} \frac{1}{1+\left(B/K_S^{B\dashv A}\right)^5}\frac{1}{1+\left(C/K_S^{C\dashv A}\right)^5} \\ \frac{1}{1+\left(C/K_S^{C\dashv B}\right)^5}\frac{1}{1+\left(A/K_S^{A\dashv B}\right)^5} \\ \frac{1}{1+\left(A/K_S^{A\dashv C}\right)^5}\frac{1}{1+\left(B/K_S^{B\dashv C}\right)^5} \end{bmatrix} \tag{1}$$

Table 1 lists the values of the parameters used in the repression interactions of all versions of the 3-gene models: the symmetric version used to generate the results of Figure 2, Figure 2–figure supplements 1 and 2, Figure 3, Figure 3–figure supplement 1 and Figure 7–figure supplement 2, the version with a weak asymmetry used in Figure 5, the version with a strong asymmetry used in Figure 5–figure supplement 1 and the version with a randomized asymmetry used in Figure 5–figure supplement 2. In the latter version, we randomly picked the values of the repression interactions of the static term $S(P)$ from a Gaussian distribution with mean $0.4$ and standard deviation $0.04$. Table 2 lists the weights $\Theta_D(g)$ and $\Theta_S(g)$ used for all 3-gene models: Models 1 and 2 used to generate the results of Figure 2, Figure 2–figure supplement 1, Figure 3, Figure 5, Figure 5–figure supplements 1 and 2 and Figure 7–figure supplement 2, as well as Models 1 and 2 with Hill functions for the weights $\Theta_D(g)$ and $\Theta_S(g)$, used in Figure 2–figure supplement 1, Figure 3–figure supplement 1 and Figure 7–figure supplement 2.

Table 1: Parameter values for the repression interactions of the 3-gene models

| Model version | $K_D^{B\dashv A}$ | $K_D^{C\dashv B}$ | $K_D^{A\dashv C}$ | $K_S^{B\dashv A}$ | $K_S^{C\dashv A}$ | $K_S^{C\dashv B}$ | $K_S^{A\dashv B}$ | $K_S^{A\dashv C}$ | $K_S^{B\dashv C}$ |
|---|---|---|---|---|---|---|---|---|---|
| Symmetric | 0.4 | 0.4 | 0.4 | 0.4 | 0.4 | 0.4 | 0.4 | 0.4 | 0.4 |
| Weak asymmetry | 0.4 | 0.4 | 0.4 | 0.36 | 0.36 | 0.4 | 0.4 | 0.4 | 0.4 |
| Strong asymmetry | 0.4 | 0.4 | 0.4 | 0.32 | 0.32 | 0.36 | 0.36 | 0.4 | 0.4 |
| Randomized asymmetry | 0.4 | 0.4 | 0.4 | 0.3825 | 0.3560 | 0.4334 | 0.4102 | 0.3802 | 0.4038 |

Table 2: Weights of the dynamic and static terms of the 3-gene models

| Weights | Model 1 | Model 2 | Model 1 with Hill functions | Model 2 with Hill functions |
|---|---|---|---|---|
| $\Theta_D(g)$ | $g^2$ | $g$ | $\frac{(g/0.4)^5}{1+(g/0.4)^5}$ | $\frac{(g/0.4)^5}{1+(g/0.4)^5}$ |
| $\Theta_S(g)$ | $(1-g)^2$ | $1-g$ | $\frac{1}{1+(g/0.6)^5}$ | $\frac{1}{1+(g/0.4)^5}$ |

###### 2.1.2 Model of Tribolium segmentation

In the model of Figure 1–figure supplement 2, the interactions between *hunchback* (*hb*), *Krüppel* (*Kr*), *mille-pattes* (*mlpt*), *giant* (*gt*) and an unidentified gene *X* are modeled (see the supplement of [1]). Note that the role of parameter $g$ is played by $caudal$ ($cad$) in the model for Tribolium segmentation.

$$P = \begin{bmatrix} hb \\ Kr \\ mlpt \\ gt \\ X \end{bmatrix} \qquad C(P) = \begin{bmatrix} -hb \\ -Kr \\ -mlpt \\ -gt \\ -X \end{bmatrix} \qquad D(P) = \begin{bmatrix} \frac{(hb/0.2)^5}{1+(hb/0.2)^5}\frac{1}{1+(Kr/0.12)^5} \\ \frac{(hb/0.4)^5}{1+(hb/0.4)^5}\frac{1}{1+(mlpt/0.25)^5}\frac{1}{1+(gt/0.01)^5} \\ \frac{(Kr/0.4)^5}{1+(Kr/0.4)^5}\frac{1}{1+(gt/0.3)^5} \\ \frac{(mlpt/0.4)^5}{1+(mlpt/0.4)^5}\frac{1}{1+(X/0.08)^5} \\ \frac{(gt/0.4)^5}{1+(gt/0.4)^5} \end{bmatrix} \tag{2}$$

$$S(P) = \begin{bmatrix} \frac{\left(hb/0.4\right)^5}{1+\left(hb/0.4\right)^5} \frac{1}{1+\left(Kr/0.4\right)^5} \\ \frac{\left(Kr/0.4\right)^5}{1+\left(Kr/0.4\right)^5} \frac{1}{1+\left(hb/0.01\right)^5} \\ \frac{\left(mlpt/0.4\right)^5}{1+\left(mlpt/0.4\right)^5} \\ \frac{\left(gt/0.4\right)^5}{1+\left(gt/0.4\right)^5} \\ \frac{\left(X/0.4\right)^5}{1+\left(X/0.4\right)^5} \end{bmatrix} \qquad \Theta_D(cad) = 3\,\frac{cad}{1+cad} \qquad \Theta_S(cad) = \frac{1}{1+cad} \tag{3}$$

#### 2.2 Gene-free models

In the gene-free model, ODEs encode flows in an abstract 2D phase space. The two geometric variables are named arbitrarily $y$ and $z$:

$$P = \begin{bmatrix} y \\ z \end{bmatrix} \qquad C(P) = \begin{bmatrix} 0 \\ 0 \end{bmatrix} \qquad D(P) = \begin{bmatrix} y\left(1-\sqrt{y^2+z^2}\right)-z \\ z\left(1-\sqrt{y^2+z^2}\right)+y \end{bmatrix} \qquad S(P) = \begin{bmatrix} \left(y_0-y\right)\left(y_1-y\right)\left(y_2-y\right) \\ -z \end{bmatrix} \tag{4}$$

where parameter $y_0$ (resp. $y_1$ and $y_2$) controls the position of the unstable fixed point (resp. the stable fixed points) of the static term along the $y$ axis. Parameter $y_0$ is set to 0 in the symmetric version of the model used in Figure 4, Figure 4–figure supplements 1, 2, 3, 4 and 5, Figure 4–movie supplements 1, 2, 3 and 4, Figure 6, Figure 7, Figure 7–figure supplement 1 and Figure 7–movie supplement 1, as well as in the asymmetric versions of Figure 6–figure supplement 1 and Figure 7–movie supplement 1. In Figure 6, parameter $y_0$ is set to 0.05 and 0.1 to model different levels of asymmetry in the basins of attraction. In Figure 7–figure supplement 1, parameter $y_0$ is set to 0.02 for Model 1 and 0.05 for Model 2 to obtain a similar level of asymmetry in the final pattern generated by the two models. We set $y_1 = -1$ and $y_2 = 1$ for all versions of the gene-free models, except for the asymmetric versions used to generate the results of Figure 6–figure supplement 1 and Figure 7–movie supplement 1. In the former, the stable fixed points of the static module are placed outside the region delimited by the limit cycle of the dynamic module by setting $y_1 = -2$ and $y_2 = 2.5$, while in the latter, we set $y_1 = -1.75$ and $y_2 = 1$.

To obtain a supercritical Hopf bifurcation with the gene-free model, we followed a similar approach than for the 3-gene model. We reasoned that the sum of the weights of the dynamic and static modules should become smaller than a degradation-like term for values of $g$ around 0.5. For this reason, an "intermediate term" $I(P) = \left[-z \quad -y\right]^T$ is introduced in the ODE. The intermediate term is weighted by the function $\Theta_I(g)$. Eq. 1 of the main text thus becomes:

$$\dot{P} = \Theta_D(g)\, D(P) + \Theta_I(g)\, I(P) + \Theta_S(g)\, S(P) + \eta(g, P) \tag{5}$$

Recall that in a given cell, only the dynamic module should be present at the beginning of the simulation, when $g = 1$. Similarly, only the static module should be present at the end of the simulation, when $g = 0$. Therefore, we set the weight of the intermediate module equal to $g\,(1-g)$, which is zero at both $g = 1$ and $g = 0$. Since this weight is of the order 2 in $g$, we make the weights of the dynamic and static modules of the order 3 in $g$ to ensure that they become smaller than the weight of the intermediate term for $g$ around 0.5. To obtain subcritical Hopf bifurcations with the gene-free model, we used a slightly different dynamic module:

$$D(P) = \begin{bmatrix} y\sqrt{y^2+z^2}\left(1-\sqrt{y^2+z^2}\right) - z \\ \\ z\sqrt{y^2+z^2}\left(1-\sqrt{y^2+z^2}\right) + y \end{bmatrix} \tag{6}$$

Table 3 lists the weights used for all gene-free models: Model 1 used to generate the results of Figure 4–figure supplements 1, 2 and 5, Figure 4–movie supplement 1, Figure 6, Figure 6–figure supplement 1, Figure 7–figure supplements 1 and 2, and Figure 7–movie supplement 1, Model 2 used to generate the results of Figure 4, Figure 4–figure supplements 1 and 5, Figure 4–movie supplement 2, Figure 6, Figure 6–figure supplement 1, Figure 7, Figure 7–figure supplements 1 and 2, and Figure 7–movie supplement 1, Model 3 used to generate the results of Figure 4–figure supplements 3 and 5, Figure 4–movie supplement 3 and Figure 7–figure supplement 2, and Model 4 used to generate the results of Figure 4–figure supplements 4 and 5, Figure 4–movie supplement 4 and Figure 7–figure supplement 2.

Table 3: Weights of the dynamic, static and intermediate terms of the gene-free models

| Weights | Models 1 and 3 | Models 2 and 4 |
|---|---|---|
| $\Theta_D(g)$ | $g^3$ | $g$ |
| $\Theta_S(g)$ | $(1-g)^3$ | $1-g$ |
| $\Theta_I(g)$ | $g\,(1-g)$ | 0 |

#### 2.3 Infinite-period scenarios of Figure 1 and Figure 7

The infinite-period scenario of Figure 1B-F is a simplified version of the model of the appendix of [4]. The dynamics of the phase of the oscillators are modeled directly using the following ODE:

$$\dot{\phi} = \omega(g) = \frac{\pi}{2}\,g^2 \tag{7}$$

The infinite-period scenario of Figure7A-E is the 1D model of coupled oscillators from [5]. In brief, the dynamics of the phase of the oscillators are described by the following ODE:

$$\dot{\phi}(x,t) = \omega(x,t) + \frac{\epsilon}{2a^2}\Big(\sin[\phi(x-a,t-\tau) - \phi(x,t)] + \sin[\phi(x+a,t-\tau) - \phi(x,t)]\Big) \tag{8}$$

where $\epsilon$ represents the coupling strength between a cell and its 2 nearest neighbors, $a$ is the average cell diameter (cd), and $\tau$ is the time delay in the coupling. The spatio-temporal profile of the frequency of the oscillators $\omega(x,t)$ is given by the following formula:

$$\omega(x,t) = \omega_\infty\Big(1 - e^{-(x-vt)/\sigma}\Big) \tag{9}$$

where $\omega_\infty$ represents the characteristic intrinsic frequency of the oscillators, $v$ is the speed at which the spatial frequency profile moves along the posterior direction, and $\sigma$ controls the spatial steepness of the frequency profile. Table 4 lists the parameter values used to generate the results of Figure 7A-E. See [5] for more details.

Table 4: Parameter values for the ODE of the phase oscillators in the infinite-period scenario of Figure 7

| $\epsilon$ [cd$^2$/min] | $a$ [cd] | $\tau$ [min] | $\omega_\infty$ [min$^{-1}$] | $v$ [cd/min] | $\sigma$ [cd] |
|---|---|---|---|---|---|
| 0.07 | 1 | 0 | 0.3886 | 0.255 | 36 |

##### 2.4 Hopf scenario of Figure 1

The Hopf scenario of Figure 1G-K is the cell-autonomous model evolved *in silico* in [6]. The model describes the dynamics of two proteins, the effector protein $E$ and the repressor protein $R$, under the control of morphogen $g$ via ODEs with time delays:

$$\dot{E} = \left(\max\left[\frac{E^{n_1}}{E^{n_1} + E_E^{n_1}}, \frac{g^{n_2}}{g^{n_2} + g_E^{n_2}}\right] \frac{S_E}{1 + (R/R_E)^{n_3}}\right)_{t-\tau_E} - \delta_E\, E \tag{10}$$

$$\dot{R} = \left(\frac{g^{n_4}}{g^{n_4} + g_R^{n_4}} \frac{S_R}{1 + (R/R_R)^{n_5}}\right)_{t-\tau_R} - \delta_R R \tag{11}$$

The subscript of a closed parenthesis indicates the time at which the expression inside the parenthesis is evaluated. If no such parenthesis with a subscript is present in a given expression, this expression is evaluated at time $t$. The values of all parameters are given in Tables 5 and 6.

Table 5: Parameter values for the ODE of the effector protein $E$ in the Hopf scenario of Figure 1

| $S_E$ | $R_E$ | $g_E$ | $E_E$ | $\tau_E$ | $\delta_E$ | $n_1$ | $n_2$ | $n_3$ |
|---|---|---|---|---|---|---|---|---|
| 0.7176 | 0.4942 | 0.0678 | 0.3213 | 0.48 | 0.8538 | 3 | 4.3549 | 4.5321 |

Table 6: Parameter values for the ODE of the repressor protein $R$ in the Hopf scenario of Figure 1

| $S_R$ | $R_R$ | $g_R$ | $\tau_R$ | $\delta_R$ | $n_4$ | $n_5$ |
|---|---|---|---|---|---|---|
| 0.9422 | 0.1156 | 0.5047 | 3.92 | 0.9759 | 3.2136 | 4.522 |

##### 2.5 Van der Pol oscillator of Figure 7

The schematic of an "asymmetric wave" profile shown on Figure 7D is generated with the following ODEs describing a Van der Pol oscillator:

$$\dot{y} = \mu\left(y - \frac{y^3}{3} - z\right) \tag{12}$$

$$\dot{z} = \frac{y}{\mu} \tag{13}$$

where parameter $\mu$ is set to 2.5. The schematic of a "symmetric wave" profile shown on Figure 7D is a sine function.

#### 3 Spatio-temporal profile of the control parameter for each model

For all models except the model for Tribolium segmentation and the infinite-period scenario of Figure 7A-E, the following function is used to describe the spatio-temporal profile of the input $g$, which is treated either as the concentration of a morphogen in the gene network models, or as an abstract control parameter in the gene-free models:

$$g(x,t) = H(x - vt) = \min\left[e^{s\,(x - vt + x_{\text{osc}})},\ 1\right] \tag{14}$$

where parameter $s$ controls the steepness of the gradient and $v$ represents the speed at which the gradient moves along the antero-posterior axis. Parameter $x_{\text{osc}}$ allows to generate a few oscillations inside the first simulated cell before $g$ starts decreasing. Note that the position vector $x$ is normalized in all our simulations, such that positions are constrained from 0 to 1. Table 7 lists the values of the parameters used for the gradients of all models (except the model for Tribolium segmentation): the gradients of the infinite-period scenario and of the Hopf scenario used to generate the results of Figure 1 B-F and Figure 1 G-K, respectively, the shallow gradient used in the 3-gene models of Figure 2, Figure 2–figure supplements 1 and 2, Figure 3, Figure 3–figure supplement 1, Figure 5, Figure 5–figure supplements 1 and 2, and Figure 7–figure supplement 2, the steep gradient used in the 3-gene models of Figure 5 and Figure 5–figure supplements 1 and 2, and the gradients used in the gene-free models of Figure 4, Figure 4–figure supplements 2, 3,

4 and 5, Figure 6, Figure 6–figure supplement 1, Figure 7, Figure 7–figure supplements 1 and 2 and Figure 7–movie supplement 1.

Table 7: Parameter values for the spatio-temporal profile of input $g$

| Model | $s$ | $v$ | $x_{\text{osc}}$ |
|---|---|---|---|
| Infinite-period scenario of Figure 1 | 0.5 | 0.08 | 0.2 |
| Hopf scenario of Figure 1 | 0.5 | 3 | 0 |
| 3-gene models, shallow gradient | 1 | 0.05 | 0.2 |
| 3-gene models, steep gradient | 2.5 | 0.05 | 0.2 |
| Gene-free models (Figure 4, its supplements and Figure 7–figure supplement 2) | 0.5 | 0.035 | 0.2 |
| Gene-free models (Figure 6 and its supplement) | 1 | 0.036 | 0 |
| Gene-free models (Figure 7 and Figure 7–figure supplement 1) | 6 | 0.0042 | 0 |
| Gene-free models (Figure 7–movie supplement 1) | 0.9 | 0.04 | 0 |

In the model for Tribolium segmentation, the role of input $g$ is played by the maternal gene $cad$. The dynamics of $cad$ is modelled with a Hill function:

$$cad(x,t) = \frac{\big(x/x^*(t)\big)^{n(t)}}{1+\big(x/x^*(t)\big)^{n(t)}} \tag{15}$$

where the time dependencies of parameters $x^*(t)$ and $n(t)$ encode respectively the regression of the morphogen gradient along the antero-posterior axis, and the gradual increase in the steepness of the morphogen gradient:

$$x^*(t) = \max\big[0.4\,,\ 0.4 + 0.2\,(t-2)\big] \qquad ; \qquad n(t) = \max\big[4\,,\ 4\,e^{(t-2)}\big] \tag{16}$$

### 4 Integration schemes

#### 4.1 Euler algorithm for deterministic simulations

Eq. 1 of the main text can be integrated via the Euler algorithm to obtain a time series representing the deterministic dynamics of vector $P$:

$$P(t+dt) = P(t) + \Bigg(\Theta_D\big(g(t)\big)\,D\big(P(t)\big) + \Theta_S\big(g(t)\big)\,S\big(P(t)\big) + C\big(P(t)\big)\Bigg)\,dt \tag{17}$$

The Euler algorithm, which is equivalent to approximating the temporal derivative of $P$ by a first-order finite difference, was used to perform deterministic simulations of all versions of the 3-gene models (Figure 2, Figure 2–figure supplements 1 and 2, Figure 5, Figure 5–figure supplements 1 and 2 and Figure 7–figure supplement 2). A similar version of this algorithm that includes the intermediate term was used for deterministic simulations of the gene-free models (Figure 4, Figure 4–figure supplements 2, 3 and 4, Figure 6, Figure 6–figure supplement 1, Figure 7, Figure 7–figure supplements 1 and 2 and Figure 7–movie supplement 1). The Euler algorithm was also used to perform simulations of the infinite-period and Hopf scenarios (Figure 1 and Figure 7). On the other hand, deterministic simulations of the model for Tribolium segmentation were carried out via the `lsoda` integrator from the `scipy` library in Python (Figure 1–figure supplement 2).

##### 4.2 Langevin equation for stochastic simulations of the 3-gene models

The stochastic nature of chemical reactions, due at least partly to the finite number of molecules involved in these reactions, introduces fluctuations in protein concentrations in single cells. To generate the results of Figure 3 and Figure 3–figure supplement 1, noise was introduced in the 3-gene models in a chemically realistic and mathematically rigorous way by following the method of [7]. In the generic formulation of the present problem, there are $N$ molecular species $S_i$, $i = 1, ..., N$, that can interact through $M$ different reactions $R_j$, $j = 1, ..., M$. Let $X_i(t)$ represent the number of $S_i$ molecules at time $t$. Then, the vector $X(t) \equiv [X_i(t) \quad ... \quad X_N(t)]$ represents the state of the whole system of $N$ molecules at time $t$. For each reaction $R_j$, a propensity function $a_j$ is defined such that if the system is in state $X$ at time $t$, then $a_j(X)\,dt$ is the probability that one $R_j$ reaction will occur in the next infinitesimal time interval $dt$, i.e. between $t$ and $t + dt$. For each reaction $R_j$, a state-change vector $\nu_j$ is defined such that its $i$th component $\nu_{ji}$ represents the change in the number of $S_i$ molecules produced by one $R_j$ reaction. Once the $M$ propensity functions and state-change vectors are defined, the time evolution of the state vector $X(t)$ is found via the $N$ deterministic reaction rate equations:

$$\dot{X}_i(t) = \sum_{j=1}^{M} \nu_{ji}\, a_j(X(t)) \qquad \text{for } i = 1\,, \dots\,, N \tag{18}$$

The numerical integration of these rate equations can be performed via the Euler algorithm:

$$X_i(t + dt) = X_i(t) + \sum_{j=1}^{M} \nu_{ji}\, a_j(X(t))\, dt \qquad \text{for } i = 1\,, \dots\,, N \tag{19}$$

The stochastic form of this simulation algorithm is given by the chemical Langevin equation:

$$X_i(t + dt) = X_i(t) + \sum_{j=1}^{M} \nu_{ji}\, a_j(X(t))\, dt + \sum_{j=1}^{M} N_j(t)\, \nu_{ji} \sqrt{a_j(X(t))\, dt} \qquad \text{for } i = 1\,, \dots\,, N \tag{20}$$

where $N_1(t)\,, \dots\,, N_M(t)$ are $M$ independent Gaussian random variables with mean and variance equal to 0 and 1, respectively, and that are not correlated in time. In the 3-gene models, the role of vector $X$ is played by $P$. Note that re-scaling the numbers of proteins $X_i$ by constant factors corresponds to multiplying both sides of Eq. 18 to 20 by that constant factor (as long as the state-change vectors $\nu_j$ are also re-scaled). Therefore, Eq. 18 to 20 are still valid when simulating protein concentrations scaled from 0 to 1 instead of absolute numbers of proteins. Furthermore, the reactions of the 3-gene models are encoded in the protein production and degradation terms. The propensities of the protein production and degradation terms are respectively $\Theta_D(g)\,D(P) + \Theta_S(g)\,S(P)$ and $P$. Eq. 19 thus becomes eq. 17, and eq. 20 can be re-written as the following expression:

$$
\begin{aligned}
P_i(t+dt) = P_i(t) &+ \Big(\Theta_D\big(g(t)\big)\,D_i\big(P(t)\big) + \Theta_S\big(g(t)\big)\,S_i\big(P(t)\big) - P_i(t)\Big)\,dt \qquad i = 1,2,3 \\
&+ \Big(N_i^{\text{prod}}(t)\,\sqrt{\Theta_D\big(g(t)\big)\,D_i\big(P(t)\big) + \Theta_S\big(g(t)\big)\,S_i\big(P(t)\big)} - N_i^{\text{deg}}(t)\,\sqrt{P_i(t)}\Big)\sqrt{dt}
\end{aligned}
\tag{21}
$$

where $N^{\text{prod}}(t) = [N_1^{\text{prod}}(t),\ N_2^{\text{prod}}(t),\ N_3^{\text{prod}}(t)]$ and $N^{\text{deg}}(t) = [N_1^{\text{deg}}(t),\ N_2^{\text{deg}}(t),\ N_3^{\text{deg}}(t)]$ are 2 vectors, each containing 3 independent Gaussian random variables with mean 0 and variance 1. This equation can be simplified by leveraging the fact that the sum of Gaussian random variables with mean 0 and different variances is equal to a single Gaussian random variable with mean 0 and a variance equal to the sum of the variances:

$$
\begin{aligned}
P_i(t+dt) = P_i(t) &+ \Big(\Theta_D\big(g(t)\big)\,D_i\big(P(t)\big) + \Theta_S\big(g(t)\big)\,S_i\big(P(t)\big) - P_i(t)\Big)\,dt \qquad i = 1,2,3 \\
&+ \Big(N_i(t)\,\sqrt{\Theta_D\big(g(t)\big)\,D_i\big(P(t)\big) + \Theta_S\big(g(t)\big)\,S_i\big(P(t)\big) + P_i(t)}\Big)\sqrt{dt}
\end{aligned}
\tag{22}
$$

where $N(t) = [N_1(t),\ N_2(t),\ N_3(t)]$ is a vector containing 3 independent Gaussian random variables with mean 0 and variance 1. Note that a different independent random variable is used for each protein, since the production term of each protein is due to a different combination of repression interactions. To control the level of noise, a parameter $\Omega$ is introduced in the previous equation such that increasing $\Omega$ decreases the level of noise:

$$
\begin{aligned}
P_i(t+dt) = P_i(t) &+ \Big(\Theta_D\big(g(t)\big)\,D_i\big(P(t)\big) + \Theta_S\big(g(t)\big)\,S_i\big(P(t)\big) - P_i(t)\Big)\,dt \qquad i = 1,2,3 \\
&+ \left(\frac{N_i(t)}{\sqrt{\Omega}}\,\sqrt{\Theta_D\big(g(t)\big)\,D_i\big(P(t)\big) + \Theta_S\big(g(t)\big)\,S_i\big(P(t)\big) + P_i(t)}\right)\sqrt{dt}
\end{aligned}
\tag{23}
$$

Since noise arises at least partly from the stochastic nature of single reactions between a finite number of proteins, increasing the concentration of proteins is expected to buffer the intrinsic chemical noise. Therefore, the noise level is expected to decrease as the protein concentration is increased. The following mathematical derivation shows that parameter $\Omega$ can be interpreted as the typical concentration of proteins in the system, such that increasing the protein concentration corresponds to increasing the value of parameter $\Omega$. First, let's take a look at the stochastic integration algorithm for protein $A$ and write explicitly the maximal production rate $\rho_A$ and the degradation rate $\delta_A$:

$$A^{+} = A + \left(\rho_A\left(\Theta_D(g)\,\frac{1}{1+\left(B/K_D^{B\dashv A}\right)^5} + \Theta_S(g)\,\frac{1}{1+\left(B/K_S^{B\dashv A}\right)^5}\frac{1}{1+\left(C/K_S^{C\dashv A}\right)^5}\right) - \delta_A\,A\right)dt$$
$$+ \frac{N_1}{\sqrt{\Omega}}\sqrt{\rho_A\left(\Theta_D(g)\,\frac{1}{1+\left(B/K_D^{B\dashv A}\right)^5} + \Theta_S(g)\,\frac{1}{1+\left(B/K_S^{B\dashv A}\right)^5}\frac{1}{1+\left(C/K_S^{C\dashv A}\right)^5}\right) + \delta_A\,A}\;\sqrt{dt} \tag{24}$$

where a + superscript on a protein concentration indicates that this variable is evaluated at time $t + dt$ and the absence of a superscript on a variable indicates that it is evaluated at time $t$. Multiplying both sides of the equation by $\Omega$ leads to the following expression:

$$\Omega\,A^{+} = \Omega\,A + \left(\Omega\,\rho_A\left(\Theta_D(g)\,\frac{1}{1+\left(B/K_D^{B\dashv A}\right)^5} + \Theta_S(g)\,\frac{1}{1+\left(B/K_S^{B\dashv A}\right)^5}\frac{1}{1+\left(C/K_S^{C\dashv A}\right)^5}\right) - \Omega\,\delta_A\,A\right)dt \tag{25}$$
$$+ N_1\sqrt{\Omega\,\rho_A\left(\Theta_D(g)\,\frac{1}{1+\left(B/K_D^{B\dashv A}\right)^5} + \Theta_S(g)\,\frac{1}{1+\left(B/K_S^{B\dashv A}\right)^5}\frac{1}{1+\left(C/K_S^{C\dashv A}\right)^5}\right) + \Omega\,\delta_A\,A}\;\sqrt{dt}$$

Now, let's re-scale all quantities that have the units of protein concentration by a factor of $\Omega$. To achieve this, we define the re-scaled variables $A^* = \Omega\,A$, $B^* = \Omega\,B$ and $C^* = \Omega\,C$, as well as re-scaled parameters $\rho_{A^*} = \Omega\,\rho_A$, $K_D^{B^*\dashv A^*} = \Omega\,K_D^{B\dashv A}$, $K_S^{B^*\dashv A^*} = \Omega\,K_D^{B\dashv A}$ and $K_S^{C^*\dashv A^*} = \Omega\,K_D^{C\dashv A}$:

$$A^{*+} = A^* + \left(\rho_{A^*}\left(\Theta_D(g)\,\frac{1}{1+\left(B^*/K_D^{B^*\dashv A^*}\right)^5} + \Theta_S(g)\,\frac{1}{1+\left(B^*/K_S^{B^*\dashv A^*}\right)^5}\frac{1}{1+\left(C^*/K_S^{C^*\dashv A^*}\right)^5}\right) - \delta_A\,A^*\right)dt \tag{26}$$
$$+ N_1\sqrt{\rho_{A^*}\left(\Theta_D(g)\,\frac{1}{1+\left(B^*/K_D^{B^*\dashv A^*}\right)^5} + \Theta_S(g)\,\frac{1}{1+\left(B^*/K_S^{B^*\dashv A^*}\right)^5}\frac{1}{1+\left(C^*/K_S^{C^*\dashv A^*}\right)^5}\right) + \delta_A\,A^*}\;\sqrt{dt}$$

A similar procedure can be followed for proteins $B$ and $C$. Therefore, multiplying the stochastic term of the Langevin equation for all proteins by $1/\sqrt{\Omega}$ is equivalent to re-scaling all variables and parameters that have the units of a protein concentration by a factor of $\Omega$. Since we set the maximal production rates and the degradation rates of all proteins to 1, the typical concentration of proteins $A$, $B$ and $C$ is normalized to 1. Re-scaling all protein concentrations and all parameters with units of protein concentration by a factor of $\Omega$ thus corresponds to setting the typical concentration of proteins to $\Omega$. In conclusion, parameter $\Omega$ of equation 23 indeed corresponds to the typical concentration of proteins.

##### 4.3 Cell-to-cell coupling in the 3-gene models

A strategy that a cell can use to fight the intrinsic noise in protein concentrations is to evaluate the protein expression state of its neighbors and change its own protein expression state accordingly. In the stochastic simulations of the 3-gene models, cell-to-cell communication is modelled via a diffusion term included in the differential equations describing the dynamics of the set of protein concentrations. The higher the concentration of a given protein is in a given simulated cell, the more this protein will diffuse to neighboring simulated cells. Diffusion thus models the process of adjusting the protein concentration of a given cell according to the protein concentration of surrounding cells. The dynamics of vector $P$ in the 3-gene models is therefore given by the following differential equation:

$$\frac{\partial P}{\partial t} = \Theta_D(g)\, D(P) + \Theta_S(g)\, S(P) - P + \eta(g, P) + D\, \frac{\partial^2 P}{\partial x^2} \tag{27}$$

where the diffusion constant $D$ controls the strength of cell-to-cell coupling. The complete stochastic simulation algorithm for the 3-gene model thus becomes:

$$\begin{aligned} P_i(x, t+dt) = P_i(x,t) &+ \left( \Theta_D\big(g(x,t)\big)\, D_i\big(P(x,t)\big) + \Theta_S\big(g(x,t)\big)\, S_i\big(P(x,t)\big) - P_i(x,t) + D\, \frac{\partial^2 P_i}{\partial x^2} \right) dt \\ &+ \left( \frac{N_i(x,t)}{\sqrt{\Omega}} \sqrt{\Theta_D\big(g(x,t)\big)\, D_i\big(P(x,t)\big) + \Theta_S\big(g(x,t)\big)\, S_i\big(P(x,t)\big) + P_i(x,t)} \right) \sqrt{dt} \end{aligned} \tag{28}$$

for $i = 1, 2, 3$. Note that diffusion is not included in the stochastic term, since diffusion of proteins is not a reaction in itself. In the simulations, the second spatial derivative is approximated by a second-order central finite difference with reflective boundaries.

##### 4.4 Stochastic simulations of the gene-free models

Since the gene-free models simulate the dynamics of abstract variables that do not represent explicitly protein concentrations, the variance of the noise is held independent of the state of the system. The stochastic algorithm used to generate the results of Figure 4–figure supplement 5 is therefore the following:

$$P_i(t+dt) = P_i(t) + \left( \Theta_D\big(g(t)\big)\, D_i\big(P(t)\big) + \Theta_I\big(g(t)\big)\, I_i\big(P(t)\big) + \Theta_S\big(g(t)\big)\, S_i\big(P(t)\big) - P_i(t) \right) dt + \frac{1}{\sqrt{\Omega}}\, N_i(t)\, \sqrt{dt} \tag{29}$$

where $i = 1, 2$, and $N(t) = [N_1(t), N_2(t)]$ is a vector containing 2 independent Gaussian random variables with mean 0 and variance 1. Parameter $\Omega$ is still included to control the level of noise, but it cannot be interpreted as the typical concentration of proteins in the system since the gene-free models do not simulate explicitly protein interactions.

#### 5 Mathematical formula for the mutual information

In deterministic simulations, the initial phase of the genetic oscillation inside a given cell determines in which part of the pattern this cell will end up. This is not necessarily the case in stochastic simulations. To quantify the robustness to noise of a given model for specific values of parameter $\Omega$ (and of the diffusion constant $D$ in the case of the 3-gene models) it is required to define a metric that measures the accuracy with which the initial phase of the genetic oscillations inside a cell predicts the region of the pattern in which this cell will end up. The specific metric used in Figure 3, Figure 3–figure supplement 1, Figure 4–figure supplement 5 and Figure 5–figure supplement 2 is the mutual information between the initial phase of the oscillator and the final protein expression state of the simulated cells. The mutual information $I(x, y)$ between two discrete variables $x$ and $y$ is given by the following expression:

$$I(x,y) = \sum_{y\in Y}\sum_{x\in X} p(x,y)\,\log\left(\frac{p(x,y)}{p(x)\,p(y)}\right) \tag{30}$$

where $X$ and $Y$ are the sets of possible values for $x$ and $y$, respectively. Intuitively, the mutual information between two variables quantifies the amount of information obtained on the value of the first variable by knowing the value of the second variable (and vice versa). If the logarithm is in base 2, the units of the mutual information are bits. To measure how precisely the phase of the oscillator is read to form the final pattern, variable $x$ is set to the phase of the oscillation in protein expression at the beginning of the simulation $\phi_i$, and variable $y$ is set to the protein expression state at the end of the simulation $P_f$:

$$I(\phi_i, P_f) = \sum_{P_f}\sum_{\phi_i} p(\phi_i, P_f)\,\log\left(\frac{p(\phi_i, P_f)}{p(\phi_i)\,p(P_f)}\right) \tag{31}$$

$$= \sum_{P_f}\sum_{\phi_i} p(P_f|\phi_i)\,p(\phi_i)\,\log\left(\frac{p(P_f|\phi_i)\,p(\phi_i)}{p(\phi_i)\,p(P_f)}\right) \tag{32}$$

$$= \sum_{P_f}\sum_{\phi_i} p(P_f|\phi_i)\,p(\phi_i)\,\log\left(\frac{p(P_f|\phi_i)}{\sum_{\phi_i} p(P_f|\phi_i)\,p(\phi_i)}\right) \tag{33}$$

To get the second equality, the fact that $p(x, y) = p(x|y)\,p(y)$ for any two variables $x$ and $y$ was used to get rid of the joint probability $p(\phi, R_i)$, which is not straightforward to evaluate directly. Similarly, the fact that $p(y) = \sum_{x\in X} p(x, y) = \sum_{x\in X} p(x|y)\,p(y)$ for any two variables $x$ and $y$ was used to get rid of $p(P_f)$, which is less easy to compute than $p(\phi_i)$. Indeed, $\phi_i$ is sampled uniformly in the simulations of the 3-gene and gene-free models, since the speed of regression of the input $g$ is constant throughout the simulations. In the 3-gene models, the different phases $\phi_i$ are defined as the different states of protein expression along the oscillation cycle generated by the dynamic module ($g = 1$). A uniform sample of $\phi_i$ is obtained by sampling this oscillation cycle at constant time intervals for a total time length of one period. In the gene-free models, the different phases $\phi_i$ are defined as the different sets of ($y$, $z$) values along the oscillation cycle generated by the dynamic module ($g = 1$). Since the oscillations are on the unit circle (centered at the origin) and have a constant speed along the cycle, sampling uniformly the angles from the positive $y$ axis (starting at 0 and stopping at $2\pi$) generates a uniform sample of $\phi_i$.

### 6 Description of the source codes

All codes are written in the `python3` programming language (except for two `Mathematica` notebooks). Commented `jupyter` notebooks can be found on Github at the following address: `https://github.com/laurentjutrasdube/Dual-Regime_Geometry_for_Embryonic_Patterning`. This repository also contains folders with the source data files, as well as the source codes used to generate the data files.

- **3-gene_det.ipynb**
  This notebook performs deterministic simulations of the symmetric 3-gene Models 1 and 2, and deterministic simulations of the 3-gene Models 1 and 2 with Hill functions for the weights of the dynamic and static modules. It also performs a bifurcation analysis of these models using the data found in the `XPPAUTO_data` folder, which also contains the `.ode` files used to generate the data with the XPP AUTO software [8]. Figure 2, Figure 2–figure supplements 1 and 2, and Figure 7–figure supplement 2 show the results obtained with this notebook.
- **3-gene_stoch.ipynb**
  This notebook performs stochastic simulations of the symmetric 3-gene Models 1 and 2, and stochastic simulations of the 3-gene Models 1 and 2 with Hill functions for the weights of the dynamic and static modules. It also generates plots of the mutual information using the data found in the `Mutual_info_data` folder, which also contains the `python` codes used to generate the data. Figure 3 and Figure 3–figure supplement 1 show the results obtained with this notebook.
- **3-gene_asym.ipynb**
  This notebook performs deterministic simulations of the asymmetric 3-gene Models 1 and 2. It also performs a bifurcation analysis of these models and generates plot of the mutual information using the data found in the `XPPAUTO_data` and `Mutual_info_data` folders, respectively. Figure 5 and Figure 5–figure supplements 1 and 2 show the results obtained with this notebook.
- **Gene-free_det.ipynb**
  This notebook performs deterministic simulations of the symmetic gene-free Models 1, 2, 3 and 4. It also performs a bifurcation analysis of these models and generates flow plots using the data found in the `XPPAUTO_data` and `Mathematica_data` folders, respectively. Figure 4, Figure 4–figure supplements 1, 2, 3 and 4, Figure 4–movie supplements 1, 2, 3 and 4, and Figure 7–figure supplement 2 show the results obtained with this notebook.
- **Gene-free_stoch.ipynb**
  This notebook performs stochastic simulations of the symmetric gene-free Models 1, 2, 3 and 4. It also generates the mutual information plots using the data found in the `Mutual_info_data` folder. Figure 4–figure supplement 5 shows the results obtained with this notebook.
- **Gene-free_asym.ipynb**
  This notebook performs deterministic simulations of the asymmetic gene-free Models 1 and 2. It also performs

a bifurcation analysis of these models using the data found in the `XPPAUTO_data` folder. Moreover, it generates plots of the flow and of the spatial wave profiles. Figure 6, Figure 6–figure supplement 1, Figure 7 and Figure 7–figure supplement 1 show the results obtained with this notebook.

- **Hopf_scenario_Fig1.ipynb**
  This notebook performs deterministic simulations of the gene network model evolved *in silico* in [6]. Results are shown on Figure 1. It also performs a bifurcation analysis of this model, shown on Figure 1–figure supplement 1.

- **Infinite-period_scenario_Fig1.ipynb**
  This notebook performs deterministic simulations of the infinite-period model of Figure 1, which is a simplified version of the model in the appendix of [4].

- **Infinite-period_scenario_Fig7.ipynb**
  This notebook performs deterministic simulations of the infinite-period model of Figure 7, which is adapted from [5].

- **Tribolium_model.ipynb**
  This notebook performs deterministic simulations of the model for Tribolium segmentation from [1]. It also generates flow plots and computes the speed of the cells in phase space. Figure 1–figure supplement 2 shows the results obtained with this notebook.
